## Supplementary figures, methods, and mathematical model for "Feedback control of organ size precision is mediated by BMP2-regulated apoptosis in the *Drosophila* eye"

T. Navarro 1, A. Iannini 1, M. Neto 1,6, A. Campoy-Lopez 1,2, J. Muñoz-García 3, P.S. Pereira 4, S. Ares 5*, F. Casares 1*
1 – CABD, CSIC/Universidad Pablo de Olavide. 41013 Seville, SPAIN.
2 – ALMIA, CABD, CSIC/Universidad Pablo de Olavide. 41013 Seville, SPAIN.
3 – Grupo Interdisciplinar de Sistemas Complejos (GISC) and Departamento de Matematicas, Universidad Carlos III de Madrid. 28911 Leganes, SPAIN.
4 – I3S, Instituto de Investigação e Inovação em Saude, Universidade do Porto; IBMC-Instituto de Biologia Molecular e Celular, Universidade do Porto, 4200-135 Porto, PORTUGAL.
5 – Grupo Interdisciplinar de Sistemas Complejos (GISC) and Centro Nacional de Biotecnologia (CNB), CSIC. 28049 Madrid, SPAIN.
6 – Present address: Instituto de Neurociencia, CSIC/Universidad Miguel Hernández, 03550 Sant Joan d'Alacant, SPAIN.

This file contains the following material:

1- SUPPLEMENTARY FIGURES and TABLES to main figures.
2- SUPPLEMENTARY STATISICAL METHODS
3- MACROS for immunostaining signal quantification
4- Mathematical model

#### SUPPLEMENTARY FIGURES AND TABLES TO MAIN FIGURES

**Suppl. Fig. 1 to Figure 1. Expression of the apoptosis markers activated caspase-3 (Cas3*) and activated Dcp-1 in control eye discs**. (a,a') *optix>GFP* primordium stained for GFP, the photoreceptor marker Elav and Cas3*. The expression of the *optix-GAL4* driver, as detected by GFP-expression is outlined (in a'). The Cas3* signal is low (a'). (b,b') *optix>+* primordium stained for the G2-marker Cyclin B (CycB) and Dcp-1. Dcp-1 signal is detected in a band of cells anterior to the differentiating wavefront (this latter lacks CycB expression).

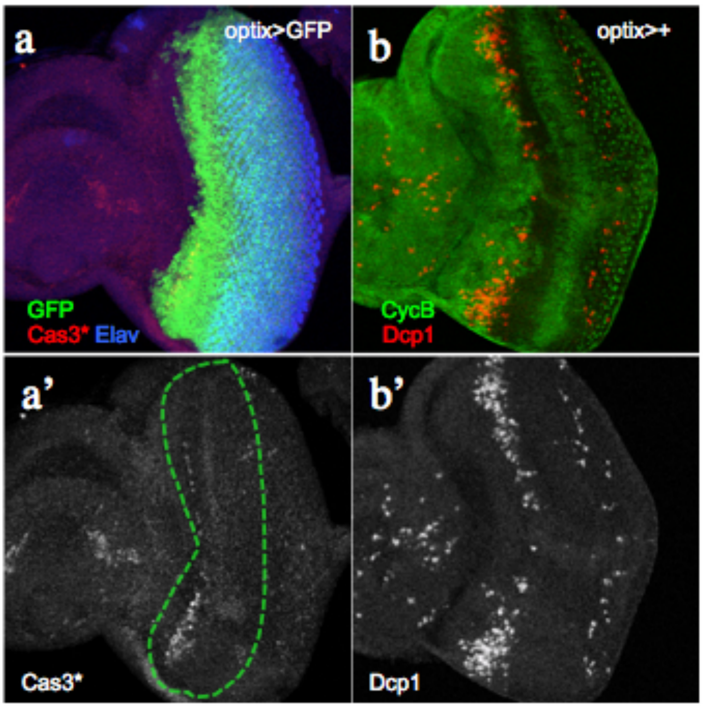


**Suppl. Table 1 to Figure 1. Analysis of the effect of genotype on rE median, rE dispersion and sFAi dispersion.** First block shows each genotype's effect on the rE median, their p.values and the predicted median for each of them. The second and third blocks show the standard deviation of each group for rE and sFAi respectively and the result of the comparison of variances of each group against the *optix>+* reference group.

| | rE | | | | | | sFAi | | |
|---|---|---|---|---|---|---|---|---|---|
| | Median effects | | | Dispersion analysis | | | Dispersion analysis | | |
| | Effects | p.value | Predicted by group (%) | sd(j) | sd(j)/sd(optix>+) | padj.(Levene test against optix>+) | sd(j) | sd(j)/sd(optix>+) | padj.(Levene test against optix>+) |
| *optix>+* | 21.11 | 0 | 21.11 | 0.62 | 1 | NA | 1.50 | 1 | NA |
| *optix>RHGRI* | 4.24 | 0 | 25.35 | 1.48 | 2.37 | 0.0015 | 2.87 | 1.91 | 0.0016 |
| *optix>tkvRI* | -3.57 | 0 | 17.54 | 1.34 | 2.14 | 0.0028 | 5.35 | 3.56 | 0 |
| *optix>tkvRI+DADRI* | 1.99 | 0 | 20.92 | 1.16 | 1.86 | 0.0028 | 2.36 | 1.57 | 0.0212 |
| *optix>DADRI* | 1.39 | 0 | 22.50 | 0.93 | 1.50 | 0.0386 | 2.84 | 1.89 | 0.0096 |
| *optix>tkvRI+RHGRI* | 1.23 | 0.0084 | 23.01 | 1.18 | 1.90 | 0.0070 | 2.90 | 1.93 | 0.0417 |
| *BAR* | -4.60 | 0 | 16.52 | 1.63 | 2.62 | 0.0001 | 5.64 | 3.75 | 0 |
| *BAR:optix>RHGRI* | 0.85 | 0.2091 | 21.61 | 1.16 | 1.87 | 0.0072 | 2.68 | 1.78 | 0.0417 |
| *BAR:optix>DADRI* | 3.26 | 0.0017 | 21.17 | 1.09 | 1.74 | 0.0070 | 2.62 | 1.74 | 0.0293 |

**Suppl. Fig. 2 to Figure 1. The average ommatidial size is approximately constant in different genotypes tested.** The average ommatidial size was calculated as the area occupied by 10 adjacent ommatidia in the equatorial region of the eye divided by 10. Independent eyes were measured on photographs of mounted eyes of the indicated genotypes. The statistical comparison of the values obtained relative to the control *("optix>+")* did not detect significant differences among genotypes.

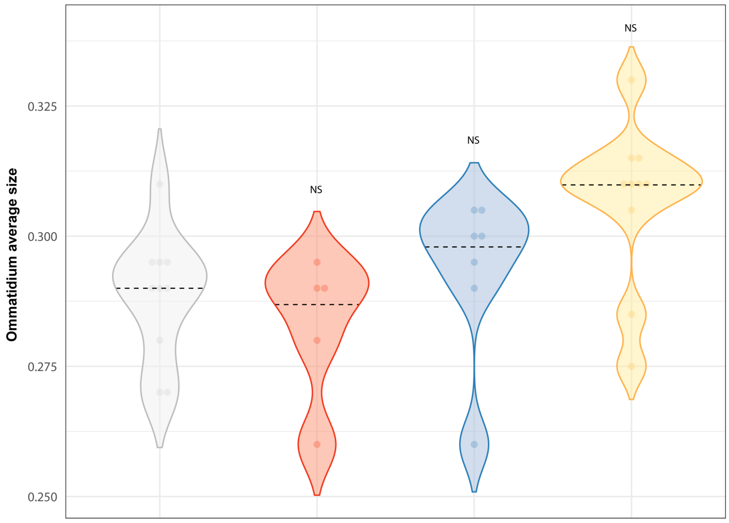


| *Median regression model* | optix>+ | optix>RHGRI | optix>tkvRI | optix>tkvRI+RHGRI |
|---|---|---|---|---|
| Effects | 0.29 | 0.00 | 0.01 | 0.01 |
| p.value | 0 | 1 | 0.2187 | 0.5596 |
| Predicted for group | 0.29 | 0.29 | 0.30 | 0.31 |

**Suppl. Fig. 3 to Figure 1. Study of the effect of a neutral RNAi on rE and sFAi values.** Results are shown for rE and sFAi of eyes from individuals of the following genotypes: "+": Oregon-R (Or-R) wild type strain; "*optix>+*": Progeny from the cross *optix-GAL4* to Or-R; "*optix>UAS-CherryRI*": The *optix-GAL4; UAS-Cherry_RNAi* progeny obtained by crossing *optix-GAL4* to *UAS-Cherry_RNAi.* This dataset was obtained independently of the datasets included in Figure 2 of the main text. The median rE of *optix>CherryRI* eyes is slightly higher than that of the *optix>+* reference control group, however, it is not significantly higher than that of the wild-type *Or-R* group (a). There are no significant differences in sFAi between any of the groups compared (b). The table shows the statistical analysis of rE and sFAi among the different groups considered. This analysis concludes that there is no significant effect of driving a non-specific, neutral RNAi on eye size precision and that the effect on final eye size is within the variability present among the alternative control lines used (An equivalent table is explained in Suppl. table 1 to Figure 2. Further details on the statistical analysis can be found in the main text Materials and Methods section).

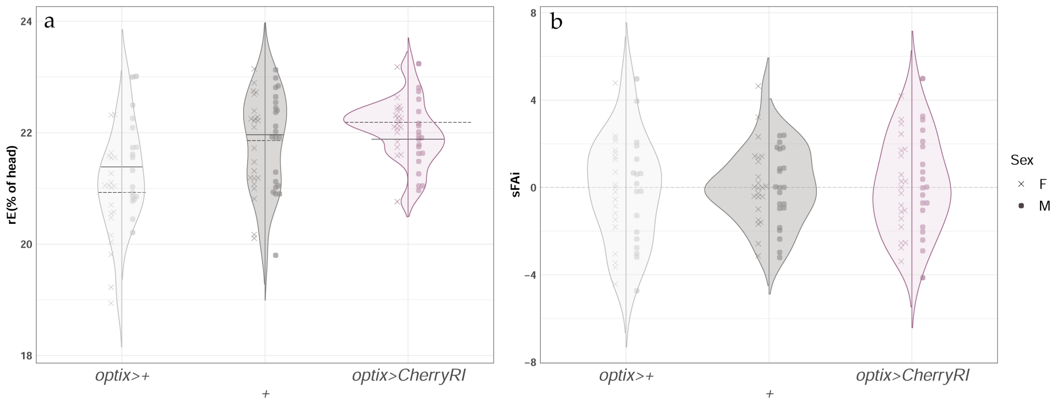


| c | rE | | | | | | sFAi | | |
|---|---|---|---|---|---|---|---|---|---|
| | Median effects | | | Dispersion analysis | | | Dispersion analysis | | |
| | Effects | p.value | Predicted by group (%) | sd(j) | sd(j)/sd(optix>+) | padj.(Levene test against optix>+) | sd(j) | sd(j)/sd(optix>+) | padj.(Levene test against optix>+) |
| *optix>+* | 21.05 | 0.0000 | 21.05 | 0.86 | 1.00 | NA | 2.41 | 1.00 | NA |
| + | 0.97 | 0.0009 | 22.02 | 0.86 | 1.00 | 0.7202 | 1.74 | 0.72 | 0.0722 |
| *optix>CherryRI* | 1.05 | 0.0000 | 22.09 | 0.54 | 0.63 | 0.0151 | 2.19 | 0.91 | 0.6170 |

**Suppl. Fig. 4 to Figure 1. Phenotypic rescue of genotypes with two UAS constructs cannot be explained by GAL4 "dilution effect".** We compared the distributions and median values of rE (a) and sFAi (b) from *optix>tkvRI* (one UAS transgene: UAS-*tkvRI*) and *optix>tkvRI +* GFP (two UAS transgenes: UAS-*tkvRI and* UAS-GFP). In the case of eye size (a), if the presence of two UAS sequences had titrated the GAL4 molecules (effectively halving the number of GAL4 molecules per UAS sequence), the expectation would be a weakening of the phenotype (derived from a weaker expression of *tkv-RNAi*). In the case of sFAi (b) titration of GAL4 would result in a reduction of the asymmetry. However, these expectations were not observed. We used the Bayes Factor (BF; shown in plot) as a measure of the strength with which a null hypothesis is accepted or rejected. Values close to 0 allow accepting the null hypothesis: "there is no GAL-4 dilution effect". Black bars show the minimum detectable differences in each comparison depending on sample sizes and distribution shapes with 80% power (See Supplementary Statistical Methods). Indeed, we find very low BF values for both rE and sFAi, supporting the idea that there is no dilution effect of GAL4 when two UAS transgenes, instead of one, are present in the genotype. Therefore, the phenotypic effects detected are genuinely caused by the genetic perturbation.

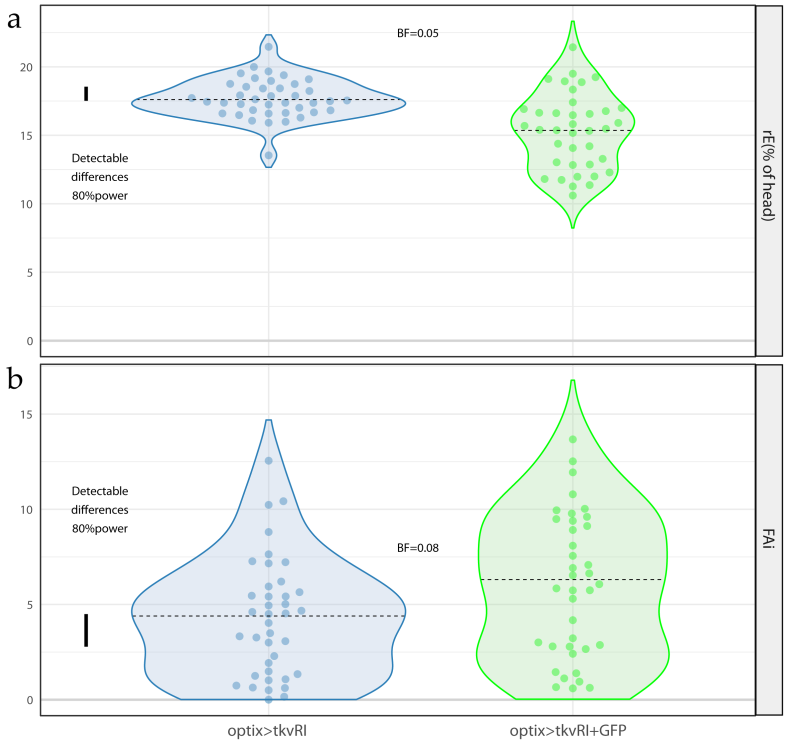


**Suppl. Figure 5 to Figure 1. Increased cell death and fragmented *dpp* expression in *Bar* discs.** L3 *dpp-Z* discs (a,b), stained for apoptosis (dcp-1, green), *dpp* transcription (beta-galactosidase, red) and the retinal differentiation marker Elav (blue). (a) control ("+": Oregon-R) and (b) *Bar* mutant discs. The ellipse approximately outlines the eye region. "*a*" marks the antennal primordium.

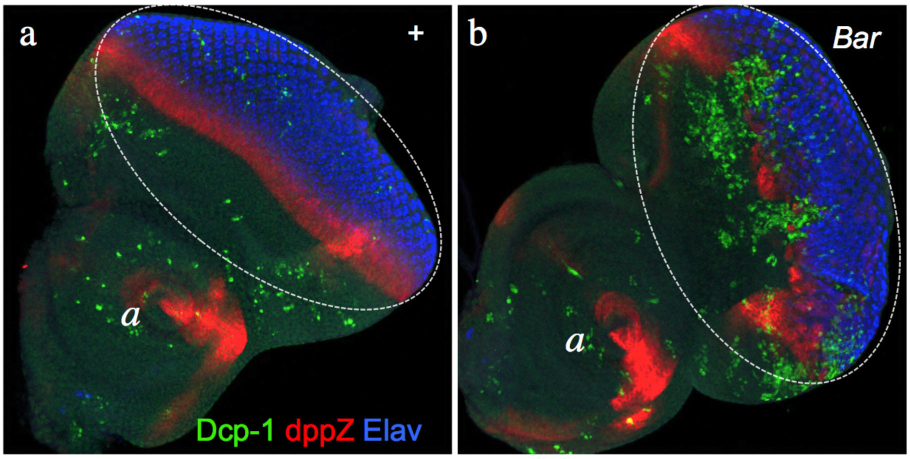

**Suppl. Fig. 6 to Figure 1. Analysis of intraindividual eye size correlation.** The intra-individual FA (FAi) is calculated using the left and right eyes *from the same individual.* The inter-individual FA (FAn) is calculated as the difference between the left eye of an individual and the right eye randomly chosen from the population of measured eyes. The difference between FAn and FAi was computed for each fly (ΔFA). If there were no "individual" effect, ΔFA should be 0. The effect of each genotype on ΔFA median, their p.values and the predicted median for each of them are shown (See Supplementary Statistical Methods). The ΔFA median is slightly, though significantly greater than zero across all genotypes relative to the control (*optix>+*). This result indicates that eye size within an individual is slightly correlated irrespective of genotype.

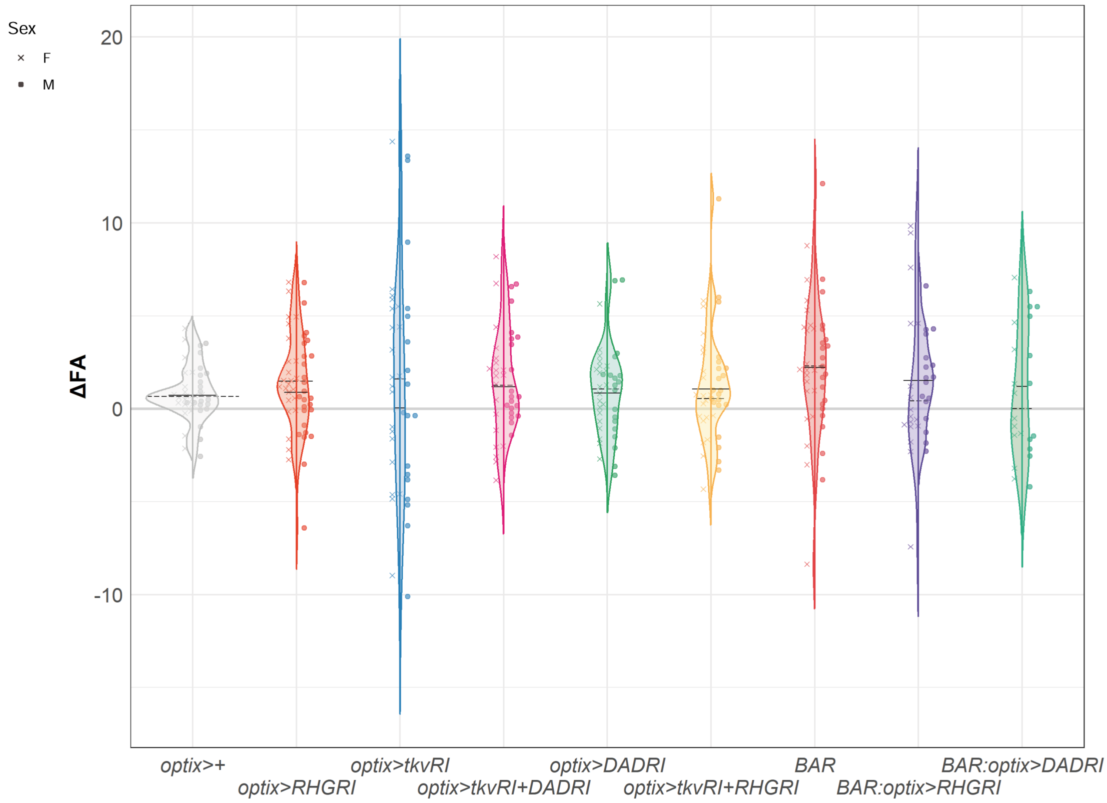


| | | | | | | | | | |
|---|---|---|---|---|---|---|---|---|---|
| Effects | 0.38 | 0.32 | 0.03 | 0.34 | 0.30 | 0.26 | 1.45 | -1.15 | -0.93 |
| p.value | 0.029 | 0.229 | 0.454 | 0.333 | 0.238 | 0.363 | 0.040 | 0.149 | 0.259 |
| Predicted by group | 0.38 | 0.71 | 0.41 | 1.34 | 0.68 | 1.00 | 1.84 | 0.72 | 1.21 |

**Suppl. Table 1 to Figure 2. Analysis of genotype effect on % of apoptotic area in the anterior domain of the eye primordium (%Apop).** Each genotype effect on %Apop median, their p.values and the predicted median for each of them are shown (See main text, Materials and Methods section).

| | optix>+ | optix>RHGRI | optix>tkvRI | optix>tkvRI+ DADRI | optix>DADRI | optix>tkvRI+ RHGRI |
|---|---|---|---|---|---|---|
| Effects | 7.21 | -5.15 | 31.21 | -25.11 | -5.25 | -32.42 |
| p.value | 0 | 0 | 0.0028 | 0.0157 | 0 | 0.0020 |
| Predicted by group (%) | 7.21 | 2.06 | 38.41 | 8.05 | 1.95 | 0.84 |

**Suppl. Fig. 1 to Figure 2. Attenuation of the Dpp signaling pathway does not affect the mitotic rate of eye progenitors.** (a, b) Representative late L3 eye imaginal discs, stained for the mitotic marker phosphor-histone H3 (PH3), and counterstained with rhodamine phalloidin (“actin”). (c) Distribution of mitotic rate, measured as the rate of PH3-positive area relative to the % of available area (non-apoptotic) in the anterior domain (regions outlined in yellow) in the two genotypes (See Supplementary Statistical Methods). Bayes Factor (BF) value close to 0 allows accepting the null hypothesis: “there is no significant differences between genotypes”. BF= 0.37. The bar indicates an 80% power to detect differences. This analysis indicates that mitotic rate of progenitors does not change in *optix>tkvRI* relative to *optix>+* controls (See Supplementary Statistical Methods).

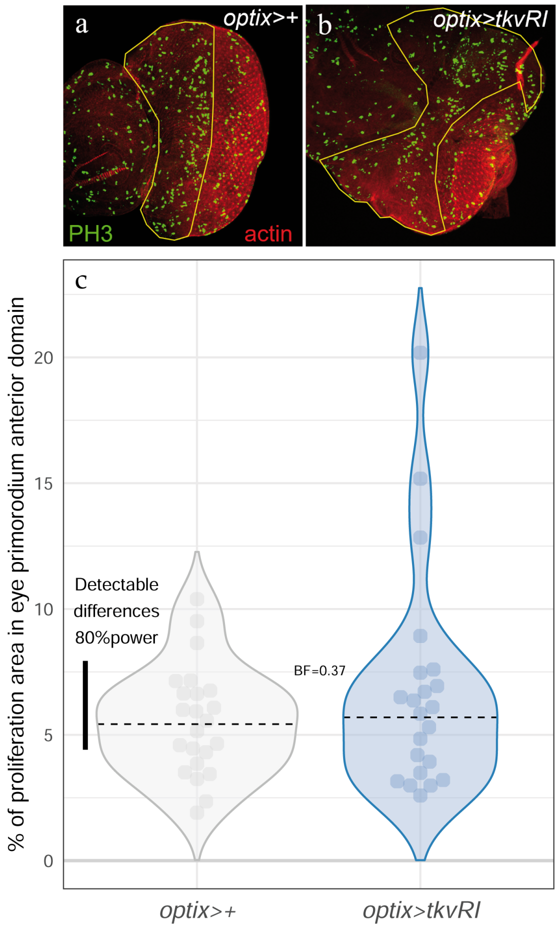

**Suppl. Fig. 2 to Figure 2. Caspase 3-associated apoptosis is induced by Dad-mediated Dpp signal attenuation.** *optix2/3-GAL4; UAS-Dad ("optix>Dad")* eye disc stained for activated caspase 3 ("cas3*"). Abundant cas3* signal is detected in cells anterior to the differentiation wave-front (dashed line). The disc is counterstained with the nuclear marker DAPI. (b,b') Close up of a cas3*-positive region. Cas3* signal (b': arrows) overlaps with pycnotic nuclei stained with DAPI (small, dense DAPI signal), indicating that cells enter an irreversible apoptotic process.

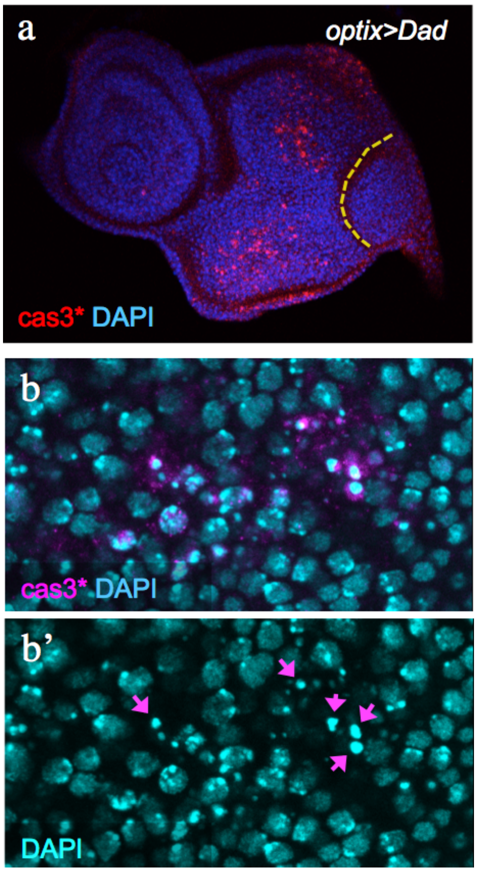

**Suppl. Fig. 1 to Figure 4. A mutant condition in which eye size varies dramatically preserving precision.** (a, b) Heads from GMR>+ (a) and GMR>Upd (b) adult males; frontal view. While the median eye size (rE: c) of *GMR>Upd* flies is approximately 1,4 times that of control (*GMR>+*) ones (statistically significant), the fluctuating asymmetry index ("precision")(FAi: d) is indistinguishable between the two genotypes. Female (F) and male (M) distributions are shown. (e, f) Eye primordia stained for the apoptotic marker Dcp-1. Apoptotic signal is observed in both *GMR>Upd* as well as in *GMR>+* control primordia.

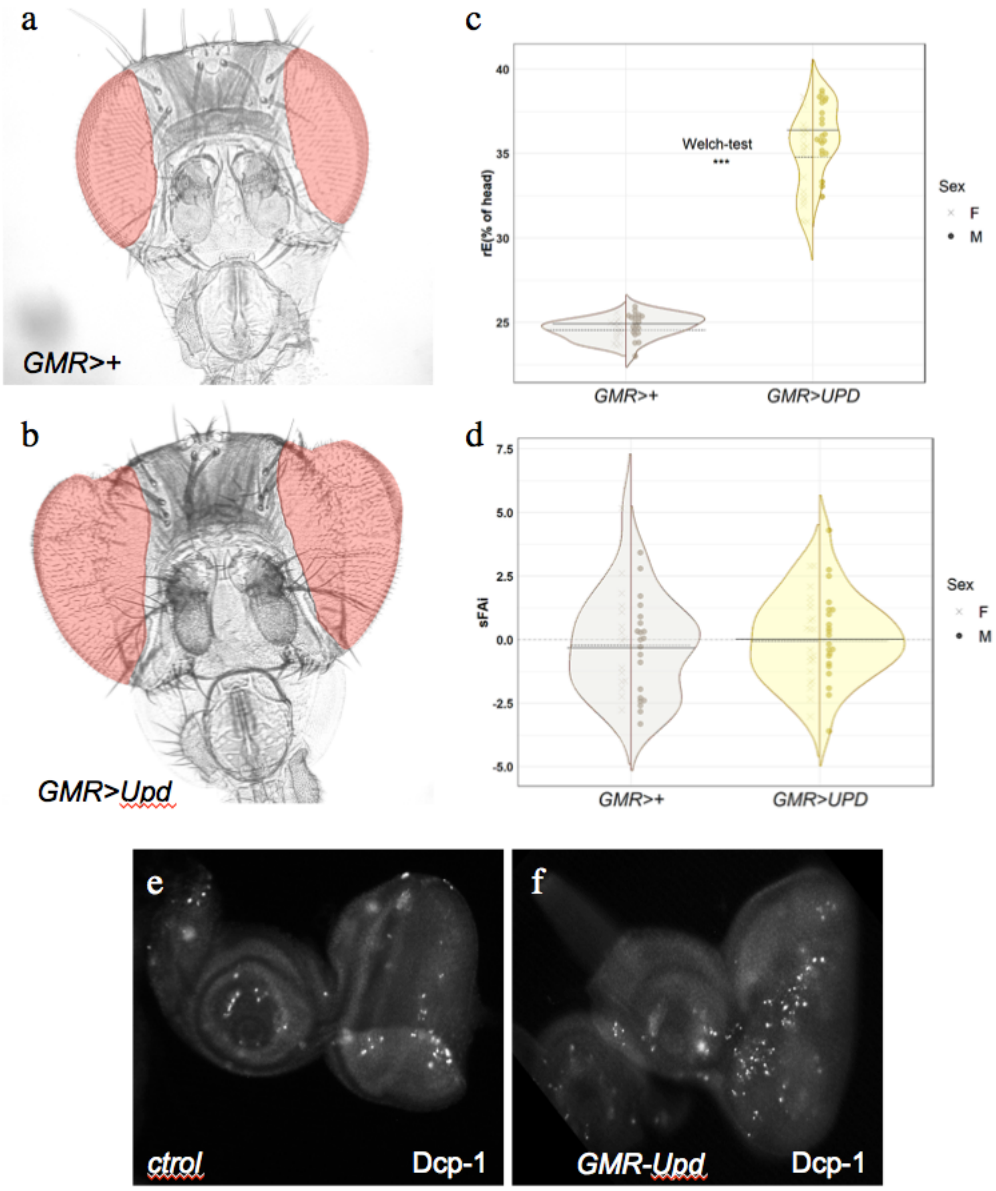

**Suppl. Figure to Materials and Methods.** Eye and head areas as measured in this work. (a,a') frontal (a) and occipital (a') focal planes, with the areas corresponding to the left eye outlined using the "lasso" tool of ImageJ. The area of the eye is the sum of both areas. (b) Area of the head, outlined using the lasso tool. The area of each eye was obtained as the sum of the area of the eye in the frontal and occipital planes (a + a').

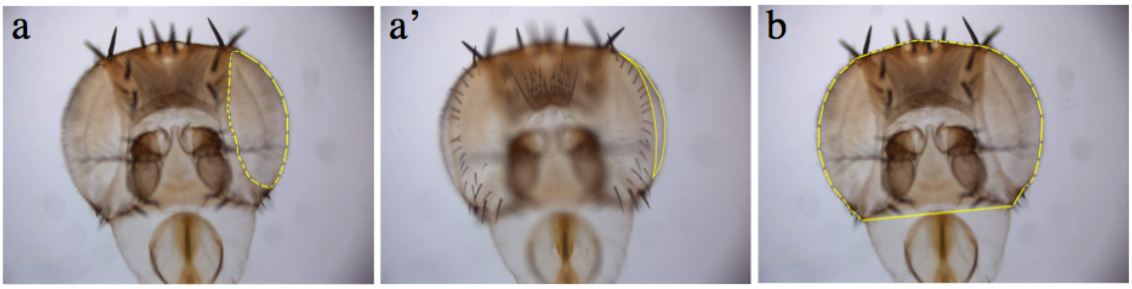


### SUPPLEMENTARY STATISTICAL METHODS

**% of proliferation area in the anterior region of the eye primordium: corrected estimation**

If a disc has a large progenitor area undergoing apoptosis, using this area to calculate the mitotic rate (%prolif) will produce an underestimation of this rate –as apoptotic cells cannot undergo mitosis. While this effect is rightly neglected in the literature for control/wild type discs (as the normal apoptotic rate is low), in our experiments, where apoptosis can be very abundant, this fact needs to be taken into account. To do so, proliferation density has to be estimated according to the proliferation-competent (non-apoptotic) anterior region area (AV). Ideally, AV could be directly calculated as the difference between total and apoptotic areas for each disc and then proliferation density estimated as a ratio. Nevertheless, measurements of proliferation and apoptosis were taken on different eye imaginal disc. The apoptotic rate (expressed as the percentage of the progenitor area expressing the Dcp-1 apoptotic marker) depends on the genotype, so we first estimated the %AV for each genotype from Dcp-1 stained discs and then applied the correction to the non-corrected raw % of proliferation (measured as the percentage of the progenitor area expressing the mitotic marker PH3), as:

$$\%\text{prolif} = \%\text{prolif(raw)}/\%\text{AV}$$

**Statistical analysis of GAL4 "dilution effect" and % of proliferation**

To test the null hypothesis that the GAL4/UAS is equally efficient in driving UAS-targets regardless of whether there is one or two UAS sequences in the genotype, we compared the phenotypes of *optix>tkvRI* (one UAS sequence) against *optix>tkvRI + GFP* (two UAS sequences). Since *optix>tkvRI* flies have smaller eyes (rE) and larger FAi than controls, if there were a "GAL4 dilution" effect (the GAL4-driven expression of the *UAS-tkv-RNAi* would be diminished in the presence of a second *UAS*), a weaker phenotype (i.e. larger rE and smaller FAi) would be expected. If $(\delta)$ expresses this dilution, we would then expect for rE and FAi that:

$$\begin{cases} H_0: \delta(\mathrm{rE}) = 0 \\ H_1: \delta(\mathrm{rE}) > 0 \end{cases}$$
$$\begin{cases} H_0: \delta(\mathrm{FAi}) = 0 \\ H_1: \delta(\mathrm{FAi}) < 0 \end{cases}$$

For the analysis of the % proliferation we do not have information on the alternative hypothesis, and the real effect could be positive or negative. Therefore:

$$\begin{cases} H_0: \delta(\%\mathrm{prolif}) = 0 \\ H_1: \delta(\%\mathrm{prolif}) \neq 0 \end{cases}$$

FAi was analyzed instead of its signed version (sFAi) in order to estimate the BF (Bayes Factor) and to determine the degree of acceptance of $H_0$ (absence of GAL4 "dilution effect" on eye size precision).

<u>Bayes factor associated with one tailed t-student test</u>

The p-value is used as a decision threshold to solve hypothesis testing from a frequentist approach. However, this decision criterion is aimed at demonstrating that $H_0$ is false but it does not, on the contrary, allow us to affirm that $H_0$ is true, since the p-value cannot be interpreted as a measure of the strength with which we accept $H_0$: not being able to demonstrate the falsity of $H_0$ does not prove its veracity. Since our aim here is precisely this, to prove $H_0$, we propose an alternative, Bayesian approach to solve the hypothesis testing. The Bayes Factor constitutes a measure of strength or confidence with which to accept $H_0$ (Dienes, 2014). Bayes factor (BF) is defined as the odds ratio between of finding the observed data under $H_1$ and under $H_0$:

$$BF = \frac{P(H_1)}{P(H_0)}$$

With values of BF greater than 1 we will tend to accept that $H_0$ is false, and true otherwise.

Whereas parameter $\delta$ distribution under the null hypothesis is well defined, under the alternative one is not, leading to a battery of possible real values for the parameter $\delta$ and subsequent density functions (Bayarri et al., 2016). ttestBF function from BayesFactor library allows to calculate BF associated to one tailed t-student test (nullInterval = c(0, Inf)).

<u>Detectable difference bars at 80% power</u>

The power of a test is fundamental in hypothesis testing, especially when aiming to prove $H_0$ (Cohen, 1988). It is defined as the probability of correctly rejecting $H_0$ and, in a complementary way, the type II error is defined as the probability of erroneously not rejecting $H_0$. We cannot confirm the veracity of $H_0$, but for a pre-specified effect size, $\delta_0$, and sample size the power of the test can be determined. If we do not know what $\delta_0$ can be expected between the 2 groups, we can proceed in the opposite way: presetting the type II error, at 20% (Cohen, 1988), and for the sample size collected calculate what would be the minimum effect size that the test could detect, $\delta_k$, so that we can judge a priori whether that capacity for discrimination is high enough to accept $H_0$ or whether the test could be missing substantial differences between groups, that is, an underpowered test.

Although the power of the test described above is classically a frequentist concept, we can transfer it to the field of Bayesian statistics if we establish a decision threshold based on the BF (Lakens, 2014) (1, as mentioned before), as is done with the p-value (typically 0.05). We followed a Montecarlo approach to estimate type II error in BF-based decision threshold for each comparison:

First, the sample empirical distributions are centered at the same mean and then shifted iteratively from 0 to the target value, $\delta_k$, the one for which the type II error of our test reaches the 20% target (80% power). Then, for each given $\delta_i$>0, a resampling of size n (number of individuals in the original samples by group and sex) from the sample empirical distributions is repeated 1000 times (*remp* function from fishmethods package) and BF calculated in each iteration (ttestBF function mentioned above). This simulation allows to calculate how many times BF was lower than 1, that is, what percentage of the times $H_0$ would be wrongly not rejected (type II error).

The target value reached, $\delta_k$, is the minimum detectable differences between groups by the BF-based decision threshold. By multiplying the resultant $\delta_k$ by the pooled standard deviation of the evaluated variable in the groups compared, we can express the minimum detectable differences in that variable's original scale and plot them as bars integrated in the sample distribution violin plots. The bar starting point is set at the *optix>tkvRI* median and oriented towards the expected value of $\delta_k$ under the alternative hypothesis (i.e.positive for rE and negative for FAi). The same method was followed to calculate detectable differences bars in the % of proliferation plot but, since we could not predict the direction of the effect in this case, the minimal detectable differences were represented both above and below the mean.

These power bars can be understood as "If real differences between populations were larger than shown by bars the test would detect them 80% of the times".

##### Significance testing of coefficients in ΔFA model

Inter-individual Fluctuation asymmetry (FAn) and Fluctuation asymmetry differences (ΔFA) were defined as follows:

$$\mathrm{FAn} = |\frac{L_i}{H_i} - \frac{R_j}{H_j}| \text{ ; } \Delta\mathrm{FA} = \mathrm{FAn} - \mathrm{FAi}$$

Where:

- i refers to the i-th individual
- j refers to random individual different from i

Due to FAn metrics (and ΔFA subsequently) for each group there are $n!^k$ possible combinations of randomized eye pairs, where n is the group size and K is the number of sex-segregated groups. This combinatorial leads to a more than $20!^{18}$ different fitted models for each of the metrics. In order to fix the problem, we used a bootstrap strategy: first, each left eye was randomly paired once and without replacement with any intragroup right eye. Then, a quantile regression model was fitted, and model coefficients were estimated (genotypic effects, See Main Statistical Methods). This resampling was repeated 1000 times. Thus, a distribution of coefficients was obtained, and 95% confidence intervals were estimated. Finally, p.values for significance testing of model coefficients were calculated also from its empirical distribution: probability of each coefficient of being smaller than 0 when coefficient estimation is positive and larger than 0 in the opposite situation.

**Real and computational results comparison**

One of the goals of this work was the implementation of a computational model of *Drosophila* eye development with which to calculate predicted eye size and precision as a function of variables regulating cellular proliferation, apoptosis and differentiation. These cellular processes can be experimentally manipulated by using targeted gene expression and attenuation methods (see Methods, main section). The computational model must yield results of eye size and precision obtained for different genotypes (dubbed here "real") after parameter fitting. The following figure shows the workflow for the validation of the computational model.

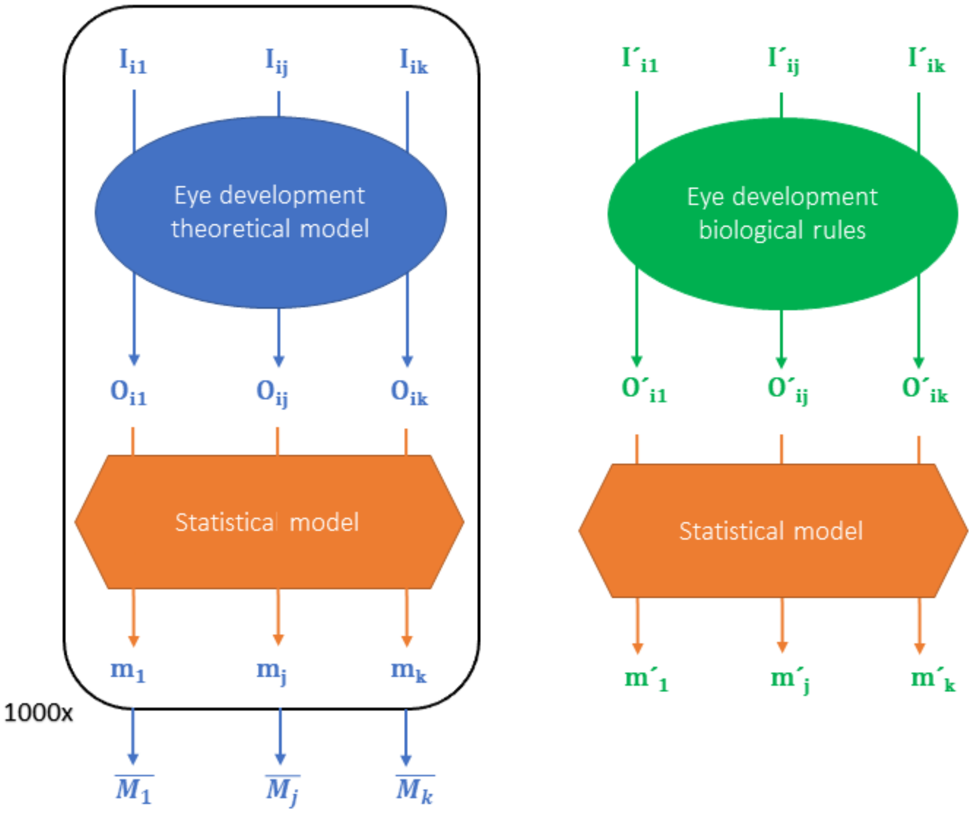


The *i*-th pair of eyes of each sample of size n is generated from the *j*-th specific combination of parameters or biological conditions (I*ij*; I´*ij*). These k conditions represent the inputs for the computational model and the biological system (the eye), respectively, and the outputs (Oij; O´ij) are values of rE and FAn. Then, the same statistical model (median regression) applied to computational and "real" (i.e. biological) data allow predicting the median of each output for each of k groups (mj;

m´j). Medians predicted by quantile regression of the output values of the computational model and biological system (mj; m´j) could be directly compared. Having into account that there is a lower limit to repeat the sampling process in the computational model we followed a Montecarlo simulation strategy by reiterating the sampling 1000 times to reach the approximated distribution of Mj.

At this point, we can pose the hypothesis to be tested in order to demonstrate that the theoretical model predicts the same eye size and precision that the respective biological genotypes:

$$\begin{cases} H_0: M'_j \sim M_j \; \forall \, j \in \; [1, \mathrm{k}] \\ H_a: \exists \, j \mid M'_j \sim \mathrm{X} \neq M_j \end{cases}$$

Under the null hypothesis at 95% confidence, the following statements must be true:

$$\mathrm{m'j} \;\in\; (\mathrm{P}_{2,5}(\mathrm{Mj}), P_{97,5}(\mathrm{Mj})) \;\forall\, \mathrm{j} \;\in\; [1, \mathrm{k}]$$

It means that the biological rE and FAn medians belong in their respective theoretical median distributions and, conversely, that the theoretical model is able to explain the observed experimental results.

Because we found a small correlation between the eyes of the same individual (Suppl. Fig. 5 to Figure 1) that cannot be accounted for by the computational model, we decided to compare the empirical FAn median, rather than the median of FAi (which is computed using left and right eye metrics of each individual) with the computational FA median distribution.

#### MACROS for immunostaining signal quantification

The following macros were designed to measure the proportion of signal of a “green-labeled” marker (in this paper PH3 or Dcp-1) to a selected are (region of interest) from confocal imaging data. They can be run with ImageJ. These are:

“Measure green channel”: Measures the green signal. It runs on all maximal projection images, as .tif files, within a folder.

“Measure green channel_threshold individual”: The same as above, but asks for a threshold signal for each image.

“Measure green channel_imagen individual”: the same as above, but for a single image.

“Genera Z projections”: Helps with bulk processing: If the data is a .lif within a folder, it allows selection of the folder and then extracts all files archives from the .lif and generates a maximal projection of each of them.

### MATHEMATICAL MODEL

#### DETERMINISTIC MODEL

We propose a model with three variables: $G$ will be a measure of the width of the progenitor population; $R_n$ a measure of the newly differentiated retinal cells, capable of producing Dpp; and $R$ a measure of the cells undergoing terminal differentiation and patterning, Fig. 1. The process will finish when all cells have differentiated and $R$ becomes a stationary value. We assume that Dpp production is proportional to $R_n$, the width of the region producing it. The effect of *hedgehog* (Hh) will be proportional to $R$. Progenitor cells can die through apoptosis, this process being repressed by Dpp. Experimental observations [1] indicate that the cell cycle becomes slower as the differentiated region of the eye becomes bigger: the doubling time of the progenitors is proportional to $R$. Denoting with $c_0$ the constant of proportionality, the rate of cell division per cell is:

$$\text{cell division rate} = \frac{\log(2)}{c_0 R}. \tag{1}$$

We will write our equations including this explicit factor $\log(2)/c_0 R$ multiplying every term; therefore this varying timescale will act as the time unit of our system, and apoptotic and differentiation rates will be expressed referred to it. In this way, our model will account for the decisions (proliferation, apoptosis, differentiation) taken by cells during the time interval corresponding to one cell cycle of the progenitors. Moreover, including the varying cell cycle duration Eq. (1) is a condition without which there is a strict experimental constraint that otherwise would be very difficult to reproduce by the model: the speed of the differentiation front is, for most of development, constant [1, 2].

Conceptually, this model can be written as a set of ordinary differential equations for the variables $G$, $R_n$ and $R$, where the evolution of each variable depends on a number of processes:

$$\frac{dG}{dt} = \frac{\log(2)}{c_0 R}(\text{proliferation} - \text{apoptosis} - \text{initial\_differentiation}) \tag{2}$$

$$\frac{dR_n}{dt} = \frac{\log(2)}{c_0 R}(\text{initial\_differentiation} - \text{final\_differentiation}) \tag{3}$$

$$\frac{dR}{dt} = \frac{\log(2)}{c_0 R}(\text{final\_differentiation}). \tag{4}$$

$G(t)$, $R_n(t)$ and $R(t)$ are functions of time $t$, but for the sake of clarity we have dropped $t$ from the notation.

Following previous models for cell proliferation [3–5], we denote the probability of a progenitor cell undergoing apoptosis along the duration of a cell cycle as the function $a(R_n)$: it is a decreasing function of $R_n$ because Dpp inhibits apoptosis. Therefore, the apoptosis term in Eq. (2) is:

$$\text{apoptosis} = a(R_n)G. \tag{5}$$

The more progenitor cells, the more new cells we have in each cycle of cell division; however, only cells not undergoing apoptosis during a cell cycle can divide. Taking this into account, the proliferation term in Eq. (2) is:

$$\text{proliferation} = (1 - a(R_n))G. \tag{6}$$

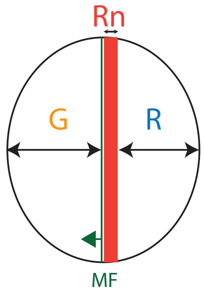


Figure 1: Schematic representation of the developing eye. To the left, progenitor cells undergo cell division, leading to the growth of the eye. In the central region, a narrow band of newly differentiated retinal cells produces Dpp. To the right, completely differentiated retinal cells will form the eye; also, they produce *hedgehog*. We denote the widths of each of these regions as: $G$ (progenitor cells), $R_n$ (newly differentiated retinal cells) and $R$ (completely differentiated retinal cells). The morphogenetic furrow (MF) roughly coincides with the frontier between progenitors and newly differentiated retinal cells, and its progression occurs at approximately constant velocity throughout most of the process.

For the rate of initial differentiation, we have to distinguish two contributions. First, it has to saturate with $G$, because beyond certain point having larger $G$ should not produce faster differentiation, since the region of $G$ differentiation is limited to the range of action of $R_n$-produced Dpp, and only cells close to the $G$-$R_n$ boundary differentiate. We model this spatial effect through a sigmoidal function $f(G)$; by imposing $f(0) = 0$ this function also ensures that when the progenitor population disappears, no new retinal cells can appear. In some sense, this limitation of the width of the progenitor cell region over which differentiation has an effect takes into account Dpp's range of action in this tissue. The second contribution to the rate of initial differentiation would be given by the activation of differentiation by *hedgehog*. We have also studied including a dependence of this rate on Dpp, but since it does not improve the results, for simplicity we do not use it. This does not exclude, however, that Dpp can play a role in this process. Hence, this contribution is an increasing function of $R$; we denote it as $h(R)$, resulting in:

$$\text{initial\_differentiation} = f(G)h(R). \tag{7}$$

Experimental observations of the width of the region of newly differentiated cells indicate that it stays almost constant throughout most of the process. We have monitored, during development, Hh signaling range, using as a read-out the active form of Ci (CiA), the nuclear transducer of the pathway. This range should be equivalent to the $R_n$ domain, as $R_n$ cells are induced by Hh signaling. The data does not show a significative trend over time in the width of this domain. This implies that cells exit the $R_n$ state at roughly the same rate that they enter it. Therefore, the rate of cells differentiating from $G \to R_n$ has to be related to the rate of cells exiting the $R_n$ state, $R_n \to R$. This can be visualized as cells in $R_n$ having some sort of *timer* that triggers final differentiation into $R$ at a certain time after becoming $R_n$. This time would correspond to the time needed for a recently recruited cell to undergo the gene expression changes required for its differentiation into a Hh-producing cell. It is not our place here to discuss the possible mechanism behind this timer, although a plausible explanation could be a roughly constant range of Hh signaling coupled to the approximately constant velocity of the differentiation wavefront. The simplest way to model this is by making the rate of final differentiation also proportional to $h(R)$:

$$\text{final\_differentiation} = b\, h(R) R_n, \tag{8}$$

where $b$ is a constant. This functional form implies that looking for a quasi-steady value of $R_n$, which is the condition $dR_n/dt \approx 0$, we would have:

$$R_n \approx f(G)/b, \tag{9}$$

which will be approximately constant as required if $f(G)$ is a saturated function of $G$, as expected during most of the process. Note that making the final differentiation rate proportional to $h(R)$ does not necessarily mean that $R$ regulates this rate; rather, it is a simplified way to model the intrinsic *timer* in $R_n$ cells that starts ticking when they undergo the initial $G \to R_n$ differentiation. When this timer runs off, $R_n$ cells differentiate into $R$, and hence the rate of this final differentiation is proportional to the initial $G \to R_n$ differentiation. Also, it should be noted that in our simplified model, having $R_n$ at a quasi-steady state and using it as a proxy for Dpp concentration means it is difficult to see the effects of Dpp purely in the dynamics of the model: that explains why we can neglect it in the initial_differentiation term, although biologically it might be a relevant factor. However, changing Dpp levels can still have a sizable effect on apoptotic probabilities.

To close the model definition, for the function modeling apoptosis we choose the Hill function:

$$a(R_n) \equiv \frac{a_0}{1 + (R_n/K)^r}, \tag{10}$$

where $a_0$ is the maximum apoptosis probability in the absence of Dpp, $K$ is the value of $R_n$ that halves the apoptosis probability, and $r$ the exponent describing the cooperativity of this repression. We choose $r = 2$ to have enough nonlinearity in the Dpp feedback. The term $a(R_n)$ is a phenomenological model of how Dpp inhibits differentiation. In a fully spatial model, the concentration of Dpp would be position-dependent, but we would still need to introduce a function of this sort. In our current model, with no space, it should be understood in a mean-field sense.

For initial differentiation, we choose to model the saturation of differentiation rate with progenitor population using the Hill function:

$$f(G) \equiv \frac{G^w}{K_G^w + G^w}, \tag{11}$$

where $K_G$ is the value of $G$ that allows the differentiation $G \to R_N$ to proceed at half its maximum potential speed, and $w$ the exponent defining the nonlinearity of the function $f(G)$. We choose $w = 1$ to ensure that for small $G$ the rate of differentiation is linear in $G$, so:

$$f(G) \equiv \frac{G}{K_G + G}. \tag{12}$$

Note that $K_G$ can be interpreted as a lengthscale of the width of the progenitor region that contributes to differentiation. Only when $G$ is of the order or smaller than $K_G$, the differentiation rate significantly decreases as the value of $G$ diminishes. This function would not be necessary in a model with explicit modeling of space and morphogen gradients. Given that we don not have data to constrain such a model, $f(G)$ is a phenomenological way of inlcuding such effects.

To model the regulation of initial differentiation by *hedgehog* we choose the function:

$$h(R) \equiv b_r + q\frac{R^u}{{K_R}^u + R^u}, \tag{13}$$

where $b_r$ is a basal differentiation rate independent of *hedgehog*, $q$ the maximum differentiation rate activated by Hh, $K_R$ the value of $R$ for which this activation is half-saturated, and $u$ the exponent defining the nonlinearity of this saturation. We choose $u = 2$ to have enough nonlinearity in the Hh feedback. The constant $b_r \neq 0$ is necessary to achieve a good fit between model and experimental data; it does not necessarily imply that differentiation can go on under the total absence of *hedgehog* or Dpp. Rather, it can model a situation where a small signal is enough to activate differentiation at a small rate, and only for higher morphogen values this rate will accelerate. The term $h(R)$ represents the effect of Hh promoting differentiation. In a spatial model Hh would be position dependent, but still a function of Hh of this sort would model the propensity of each cell to differentiate. As with $a(R_n)$, the function $h(R)$ has to be interpreted as a mean-field version of a spatially extended $h(Hh(x,y))$. Therefore, the constant $K_r$ is not modeling a lengthscale, but a threshold of Hh activity. To maintain dimensional consistency with $R$, which itself has units of space, $K_r$ is also expressed in terms of spatial dimensions. However, please note that $K_r$ actually signifies the concentration of Hh at which its effect on differentiation reaches half saturation. Remember that we are taking $R$ as a proxy for Hh levels. If we were to replace $R$ by a proportional Hh concentration, the same proportionality constant would apply to $K_r$ and convert it to its true dimension as a concentration. Therefore, contrary to $K_G$, the value of the phenomenological constant $K_R$ should not be considered a meaningful lengthscale.

The regulatory logic implied by these definitions is shown in Fig. 2, and allows us to write the qualitative Eqs. (2)-(4) as the quantitative equations:

$$\frac{dG}{dt} = \frac{\log(2)}{c_0 R}((1 - a(R_n))G - a(R_n)G - f(G)h(R)) \tag{14}$$

$$\frac{dR_n}{dt} = \frac{\log(2)}{c_0 R}(f(G)h(R) - b\,h(R)R_n) \tag{15}$$

$$\frac{dR}{dt} = \frac{\log(2)}{c_0 R} b\,h(R)R_n. \tag{16}$$

Substituting the expressions for $a(R_n)$, $f(G)$ and $h(R)$, we have:

$$\frac{dG}{dt} = \frac{\log(2)}{c_0 R}\left(\left(1 - \frac{2a_0}{1 + (R_n/K)^2}\right)G - \left(b_r + q\frac{R^2}{{K_R}^2 + R^2}\right)\frac{G}{K_G + G}\right) \tag{17}$$

$$\frac{dR_n}{dt} = \frac{\log(2)}{c_0 R}\left(b_r + q\frac{R^2}{{K_R}^2 + R^2}\right)\left(\frac{G}{K_G + G} - b\,R_n\right) \tag{18}$$

$$\frac{dR}{dt} = \frac{\log(2)}{c_0 R}\left(b_r + q\frac{R^2}{{K_R}^2 + R^2}\right) b\,R_n. \tag{19}$$

As specified, these equations are deterministic. Next we will work out the way to include the effects of noise and fluctuations.

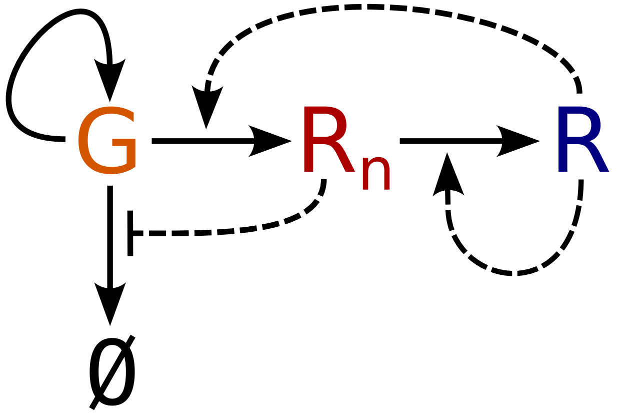


Figure 2: Regulatory logic of the model. Continuous arrows denote cellular transitions, dashed lines are positive (pointed arrow) or negative (blunt arrow) regulations of these transitions. Progenitor cells, $G$, can proliferate, differentiate into newly differentiated retinal cells, $R_n$, or die through apoptosis (∅). Dpp produced by $R_n$ inhibits apoptosis, *hedgehog* produced by finally differentiated retinal cells, $R$, promotes differentiation. $R_n$ cells maturate into $R$ cells.

**PARAMETER VALUES**

We have fitted parameter values to reproduce the dynamics of eye growth in *optix*>+, Figure 5 of the Main Text. Although Figure 5 does not show data for $R_n$, we have made independent observations of its typical value that have also been used to constrain the model. The parameter definitions and values are listed in the table below.

Table 1: Definition of the parameters in the model and values used for *optix*>+. Non-dimensional rate constants indicate proportionality factors to the cell division rate $\log(2)/c_0 R$.

| | | |
|---|---|---|
| $c_0$ | progenitor's doubling time is $c_0 R$ | 0.1 h $\mu$m$^{-1}$ |
| $a_0$ | maximum apoptosis probability | 0.3 |
| $K$ | value of $R_n$ that halves the apoptosis probability | 4.5 $\mu$m |
| $b_r$ | basal differentiation rate independent of Hh | 18 |
| $q$ | maximum differentiation rate activated by Hh | 670 |
| $K_R$ | value of $R$ for which Hh activation is half-saturated | 190 $\mu$m |
| $K_G$ | progenitor lengthscale contributing to differentiation | 4.0 $\mu$m |
| $b$ | final differentiation rate constant | 0.04 |

To reproduce the experimental data in Figure 5 of the Main Tex, the initial condition we used for all simulations was $G = 32\mu$m, $R_n = 10\mu$m, $R = 5\mu$m. We have checked that our results are robust to adding reasonable amounts of noise to these initial conditions; moreover, the effects caused by this variability are difficult to distinguish from those of dynamic stochasticity (see below). Therefore, for simplicity, we do not discuss these observations.

The data we used to determine the parameter set, corresponding to a single condition, cannot univocally determine each individual parameter. This is a general characteristic of biological and physical models, termed *sloppiness* [6–8]. This is not a problem while not trying to reach conclusions from individual parameter values: an equivalent fit with a different parameter set produces the same results in the range of validity of the model. We have indeed checked that our results are independent of the specific parameter set in the table above. In this sense, the parametrized deterministic model is a tool to understand the changes to it that lead to different conditions, and the mechanisms of modeling noise that reproduce the fluctuations observed experimentally.

**MODELING RNAi TREATMENTS**

We blocked apoptosis by driving, specifically anterior to the differentiation wavefront, a polycistronic RNAi against *reaper*, *hid* and *grim* (RHG), the three major pro-apoptotic genes in *Drosophila*: *optix*>*RHGRI* (see experimental Materials and Methods for details). The reduced apoptosis in *optix*>*RHGRI* is modeled using a lower value of $a_0$, the maximum apoptosis probability.

The Dpp pathway can be specifically attenuated in eye progenitors during development, by targeting the expression of an RNAi against the Dpp receptor *tkv* (thickveins) in these cells (*optix*>*tkvRI*). To confirm that results on *optix*>*tkvRI* result from the attenuation of the Dpp pathway, we drove expression of RNAis against *dad* (*daughters against-dpp*), a Smad molecule that acts as a feedback inhibitor of the pathway (*optix*>*DADRI*). Simultaneous use of both RNAis results in *optix*>*tkvRI+DADRI*.

To model these treatments we modify the parameter $K$, that sets the sensibility of apoptosis to the value of $R_n$: for *optix*>*tkvRI* we increase $K$, reducing the capability of $R_n$ to inhibit apoptosis; for *optix*>*DADRI* we decrease $K$, making lower $R_n$ values to have a stronger effect inhibiting apoptosis. Since we do not study serial treatments with different concentrations of tkvRI and DADRI, modeling with functions of their concentrations is unjustified and cannot be constrained. Moreover, since the effects of tkvRI and DADRI are surely nonlinear, a factor modeling the combination *optix*>*tkvRI+DADRI* is not necessarily a simple function of the factors $tkvRI$ and $DADRI$. Therefore, we also vary $K$ by a factor that in principle can be greater or smaller than 1.

We also performed experiments combining RNAis targeting *RHG* and *tkv*: *optix*>*tkvRI+RHGRI*. Again, the combination of these two treatments may lead to non-trivial effects; in particular, the reduction of Dpp feedback by RNAi targeting *tkv* may lead to a reduction of the anti-apoptotic effect of the RNAi against RHG. Therefore, to model *optix*>*tkvRI+RHGRI* we change the parameter $K$ by the same factor as in *optix*>*tkvRI*, but allow for a lower reduction of $a_0$ than in the *optix*>*RHGRI* case.

In the following table, we list the parameters used to model the effects on apoptosis of these different conditions:

Table 2: Parameters used to simulate different conditions.

| condition | parameter | value |
|---|---|---|
| *optix*>*RHGRI* | $a_0$ | 0.011 |
| *optix*>*tkvRI* | $K$ | 6.8$\mu$m |
| *optix*>*DADRI* | $K$ | 3.7$\mu$m |
| *optix*>*tkvRI+DADRI* | $K$ | 4.6$\mu$m |
| *optix*>*tkvRI+RHGRI* | $K$ | 6.8$\mu$m |
| *optix*>*tkvRI+RHGRI* | $a_0$ | 0.077 |

#### LINEAR STABILITY OF THE DETERMINISTIC MODEL

To understand the effects of the regulatory logic in Fig. 2, we perform a linear stability analysis of the deterministic model given by Eqs. (14)-(16). Only some generic properties of the functions $a(R_n)$, $f(G)$ and $h(R)$ will be necessary, so the conclusions of this analysis do not depend on the specific choices we have made for these functions. First we have to determine the fixed points of the dynamics of Eqs. (14)-(16), these are the points for which:

$$\frac{dG}{dt} = 0 \tag{20}$$

$$\frac{dR_n}{dt} = 0 \tag{21}$$

$$\frac{dR}{dt} = 0. \tag{22}$$

Assuming the Hh feedback $h(R)$ as a strictly positive function implies that necessarily any fixed point of the deterministic Eqs. (14)-(16) has to fulfill $R_n = 0$. In turn, given $a(R_n) < 1$ (as corresponds to a probability) and that $f(G)$ is a positive function with $f(0) = 0$ (there can be no initial differentiation when there are no progenitors), the other condition for a fixed point is $G = 0$. The variable $R$ is not determined uniquely determined at the fixed point: it depends on the initial conditions. We denote the value of $R$ when $G = R_n = 0$ as $R^*$, and it is always unstable, since any positive value of $R_n$ will make it change. However, if we can ensure stability of the $G = 0$, $R_n = 0$ fixed point, this will not be a problem, since this fixed point is an *absorbing state*: once the $G$ and $R_n$ pools are depleted, there are no perturbations that can make new cells of these populations appear. So we only have to study the stability with respect to $G$ and $R_n$, Eq. (14) and Eq. (15). For our choice of an initial differentiation rate that is a linear function of $G$ for small $G$, the Jacobian of Eqs. (14–15) evaluated at the fixed point $G = 0$, $R_n = 0$ and $R = R^*$ is:

$$J = \begin{pmatrix} \frac{\partial \dot{G}}{\partial G} & \frac{\partial \dot{G}}{\partial R_n} \\ \frac{\partial \dot{R}_n}{\partial G} & \frac{\partial \dot{R}_n}{\partial R_n} \end{pmatrix} = \frac{\log(2)}{c_0 R^*} \begin{pmatrix} 1 - 2a(0) - f'(0)h(R^*) & 0 \\ f'(0)h(R^*) & -bh(R^*) \end{pmatrix}, \tag{23}$$

where $f'(0)$ is $\partial f / \partial G$ evaluated at $G = 0$. The eigenvalues of $J$ are its diagonal entries,

$$\lambda_1 = 1 - 2a(0) - f'(0)h(R^*) \tag{24}$$

$$\lambda_2 = -bh(R^*). \tag{25}$$

The condition for stability of the fixed point is that these eigenvalues are negative. For a positive function $h(R)$, $\lambda_2$ automatically fulfills the condition; from $\lambda_1$ we obtain the stability condition:

$$2a(0) + f'(0)h(R^*) > 1. \tag{26}$$

It can be readily seen that increasing values of $a(0)$, $f'(0)$ and $h(R^*)$ favor stability: higher apoptosis probability for low Dpp, faster rate of initial differentiation and strong Hh feedback favoring initial differentiation contribute to the stability of the fixed point.

For our specific choices of $a(R_n)$, $f(G)$ and $h(R)$ in Eqs. (17–18), we have $a(0) = a_0$ and $f'(0) = 1/K_G$, yielding the Jacobian:

$$J = \frac{\log(2)}{c_0 R^*} \begin{pmatrix} 1 - 2a_0 - \left(b_r + q\frac{R^{*2}}{{K_R}^2 + R^{*2}}\right)/K_G & 0 \\ \left(b_r + q\frac{R^{*2}}{{K_R}^2 + R^{*2}}\right)/K_G & -b\left(b_r + q\frac{R^{*2}}{{K_R}^2 + R^{*2}}\right) \end{pmatrix}, \tag{27}$$

The stability condition now reads:

$$2a_0 + \left(b_r + q\frac{R^{*2}}{{K_R}^2 + R^{*2}}\right)/K_G > 1, \tag{28}$$

cleanly displaying that Dpp regulates stability through apoptosis, and Hh through differentiation. Experimental measurements of the apoptotic area in differentiated eyes show that it is around 40% in conditions in which apoptosis is not repressed. This would imply a value of $a_0 \approx 0.4$, meaning that the possibility of apoptosis, just by itself, is close to be enough to ensure stability of the fixed point, since $2a_0 \approx 0.8$ is close to $1$.

**STOCHASTIC MODEL**

In order to introduce the effect of fluctuations, it will be convenient to rewrite the model in Eqs. (14)-(16) using a vector notation. Defining $X(t)$ as the vector

$$X(t) = \begin{pmatrix} G(t) \\ R_n(t) \\ R(t) \end{pmatrix}, \tag{29}$$

where $t$ is time, we can write:

$$dX = F(X(t))dt, \tag{30}$$

where $F(X(t))$ is the so-called drift vector:

$$F(X) = \frac{\log(2)}{c_0 R}\begin{pmatrix} (1 - a(R_n))G - a(R_n)G - f(G)h(R) \\ f(G)h(R) - b\,h(R)R_n \\ b\,h(R)R_n \end{pmatrix}. \tag{31}$$

A standard way to include fluctuations in this kind of model, interpreting the different transition rates in a sense similar to propensities of chemical reactions, is to formulate the model as a system of Langevin equations [9,10]. Langevin equations are a type of stochastic differential equation (SDE) that, under certain conditions, are reasonable approximations of the stochastic dynamics of a set of chemical reactions or population. The main microscopic limits of this approximation translate, at an observational level, in the fact that the Langevin approximation breaks down when the noise distributions are multimodal [11]. In our work, the experimental results show noise distributions with a single mode, Figures 1 and 4 of the main text, therefore allowing the use of Langevin dynamics to model them. The noise term in an SDE can be interpreted mathematically in two different ways: the Itô or the Stratonovich sense. The distinction is very technical and out of the scope of our discussion: in our case, the noise has to be interpreted in the Itô sense [9,10]. The Langevin equation associated to the model in Eq. (30) is:

$$dX = F(X(t))dt + M(X(t))dW \tag{32}$$

where $W$ is a vector of the four independent Wiener processes [9], representing the fluctuations associated to the four kind of processes in the model (proliferation, apoptosis, initial differentiation and final differentiation), and $M(X(t))$ is the diffusion matrix encoding the different contributions to the noise. This noise that originates from the stochasticity of birth, death, and interaction processes in a population is usually termed *demographic noise*.

The diffusion matrix corresponding to the model with drift vector given by Eq. (31) is [10]:

$$M(X) = \sqrt{\frac{\log(2)}{c_0 R}} \begin{pmatrix} \sqrt{\frac{(1-a(R_n))G}{V_p}} & \sqrt{\frac{a(R_n)G}{V_a}} & -\sqrt{\frac{f(G)h(R)}{V_i}} & 0 \\ 0 & 0 & \sqrt{\frac{f(G)h(R)}{V_i}} & -\sqrt{\frac{b\,h(R)R_n}{V_f}} \\ 0 & 0 & 0 & \sqrt{\frac{b\,h(R)R_n}{V_f}} \end{pmatrix}, \tag{33}$$

where the $V$ parameters determine the noise strength associated to each process: $V_p$ is proportional to the inverse of the noise strength related to proliferation, $V_a$ to apoptosis, $V_i$ to the initial differentiation $G \to R_n$ and $V_f$ to the final differentiation $R_n \to R$. Note that this matrix is non-diagonal: the fluctuations of different variables (rows of the matrix) that come from the same processes (columns of the matrix) are related. The random fluctuations are encoded in the stochastic vector $dW$ that multiplies this matrix, $M(X)$ modulates those random numbers to produce the effect of the noise on each variable. Considering:

$$dW = \begin{pmatrix} dW_p \\ dW_a \\ dW_i \\ dW_f \end{pmatrix}, \tag{34}$$

as the *random rolls of dice* at each infinitesimal of time $dt$ for the processes of proliferation, apoptosis, initial differentiation, and final differentiation, respectively, the vector $M(X)dW$ gives us the stochastic part of the equations of each of our three variables $G$, $R_n$ and $R$:

$$M(X)dW = \sqrt{\frac{\log(2)}{c_0 R}} \begin{pmatrix} \sqrt{\frac{(1-a(R_n))G}{V_p}}dW_p + \sqrt{\frac{a(R_n)G}{V_a}}dW_a - \sqrt{\frac{f(G)h(R)}{V_i}}dW_i \\ \sqrt{\frac{f(G)h(R)}{V_i}}dW_i - \sqrt{\frac{b\,h(R)R_n}{V_f}}dW_f \\ \sqrt{\frac{b\,h(R)R_n}{V_f}}dW_f \end{pmatrix},$$

Note the opposite sign of the expressions multiplying $dW_i$: a fluctuation in initial differentiation affects in the same quantity but opposite sign the dynamics of $G$ and $R_n$. The same applies for fluctuations associated to final differentiation, $dW_f$: they have the same value but opposite sign for $R_n$ and $R$.

Stochastic equations have been numerically integrated using the Euler-Maruyama method from the function sde_euler in SDETools, a Matlab toolbox for the numerical solution of stochastic differential equations [12]. The option 'NonNegative' is set to 'yes', to ensure that the noise does not drive to negative values variables that are positive by definition.

Our model uses continuous variables, as opposed to the discrete nature of cells. To avoid that fluctuations unrealistically keep growth going when the values of $G$ or $R_n$ are well below the size of a cell, we set a threshold of $0.1\mu$m, below which we make these variables zero. The multiplicative character of noise ensures that then fluctuations also disappear.

For the $V$ parameters determining noise strength, we have used the values in the following table:

Table 3: Parameters used to define the noise strength associated to each process. For *optix>+* we have multiplied these values by a factor 6, making the noise weaker.

| | | |
|---|---|---|
| $V_p$ | inverse of the noise strength related to proliferation | 100 |
| $V_a$ | inverse of the noise strength related to apoptosis | 1.0 |
| $V_i$ | inverse of the noise strength related to initial differentiation | 6000 |
| $V_f$ | inverse of the noise strength related to final differentiation | 6000 |

The exact value of the parameters is not that relevant: the key observation is that, to reproduce the high asymmetry observed between eyes in *optix>tkvRI*, apoptosis has to be a process much noisier than all others. From an intuitive point of view, this makes sense: differentiation is driven by well organized morphogens, but apoptosis in our model cells is just a probability for each cell to die randomly during a cell cycle.

Figure 6 of the Main Text shows that our model does an excellent job reproducing the eye sizes for different conditions. The fit of the FAi in different conditions could also have been almost perfectly good if we had allowed for different values of the noise parameters in each condition. We have chosen not to do so and not to fit the FAi with different $V$ parameters for different conditions; instead, we use a single parameter set for the noise and let the changes in apoptotic mechanics in each condition to be the sole drivers of differences in asymmetry. However, conditions like *optix>tkvRI+DADRI* rescue the eye size but present higher asymmetry values than *optix>+*. The *optix>RHGRI* condition, that we have modeled as a reduction in the probability of apoptosis, is also more noisy than *optix>+*. This is difficult to reconcile with the observation, Figure 6 of the Main Text, that *optix>+* presents lower FAi than any of these conditions. Since second-order effects of these conditions on the noise are out of the scope of our model, for simplicity to approximate the FAi for *optix>+* we have multiplied all the $V$ parameters in Table 3 by a factor $6$, therefore reducing the noise for this condition.

**NOISE STRENGTH OF THE DIFFERENT PROCESSES**

Our determination of the parameter values of the noise strength associated with each process, Table 3, shows that of all the processes included in the model, apoptosis has to be the noisiest ($V_a$ is small compared to other noise parameters) in order to explain the high asymmetry between eyes observed when apoptosis is not repressed. To illustrate how this is indeed the case, we show the comparison of simulation and experimental results for eye size and asymmetry, Figure 3. We have then studied the effect of having weaker noise associated with apoptosis, Figure 4: in this case FAi obtained from simulations for *optix>tkvRI* is similar to other conditions. Strong noises in the other processes while keeping apoptotic noise low, Figures 5–7, produce high FAi for all conditions, not discriminating *optix>tkvRI* from the other conditions. From the nature of the conditions we have studied, where apoptosis is primarily affected, we expected that the high asymmetry of the *optix>tkvRI* condition would be associated with apoptotic noise, while a strong contribution to noise of other processes would not discriminate this condition and produce high levels of asymmetry in all the conditions we have simulated. Our simulations support this picture.

For a more systematic understanding of noise effects, we have calculated the Fluctuating

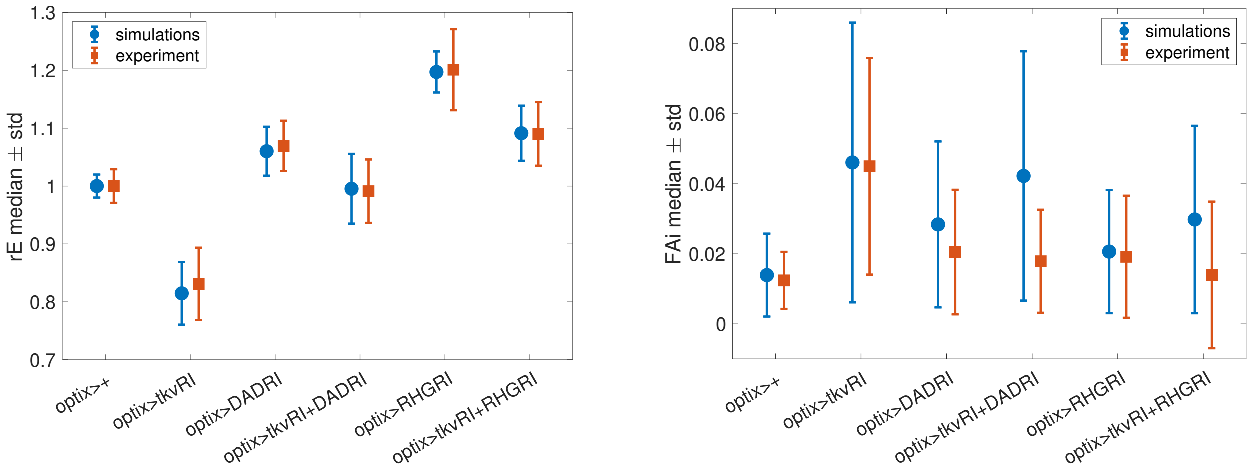


Figure 3: Standard noise (strong apoptotic noise): comparison for different conditions, as indicated, of simulation and experimental results. Simulations of 100 eyes with parameters in Tables 1–3. Left: eyes size, normalized to allow comparison of simulation and experiment. $rE$ is the average intraindividual eye size. Right: eyes asymmetry. $FAi$ is the intraindividual factor of asymmetry. Virtual *flies* with two eyes have been created by making all possible pairs of the 100 eyes.

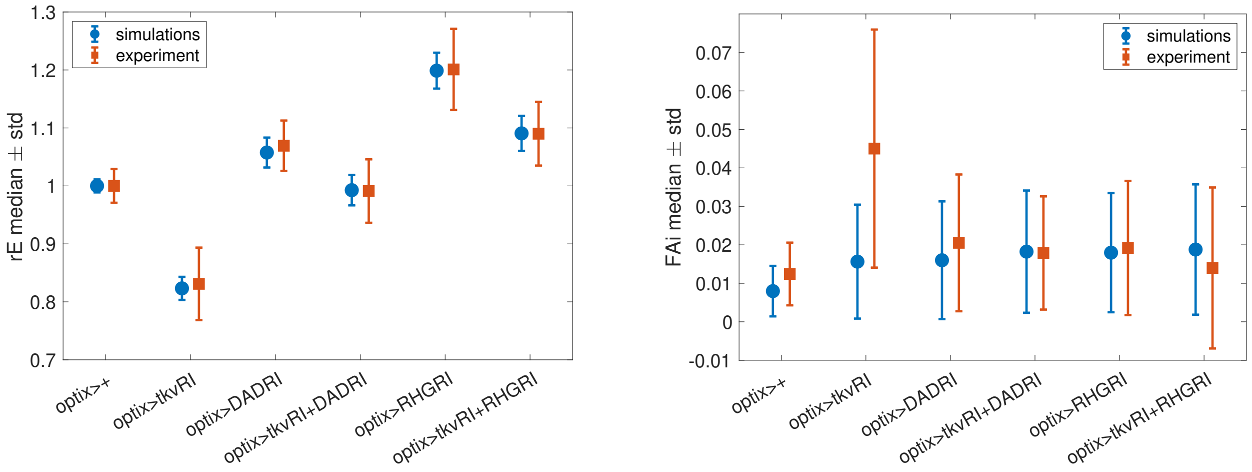


Figure 4: Weak apoptotic noise: comparison for different conditions, as indicated, of simulation and experimental results. Simulations of 100 eyes with $V_a = 100$, other parameters as in Tables 1–3. Left: eyes size, normalized to allow comparison of simulation and experiment. $rE$ is the average intraindividual eye size. Right: eyes asymmetry. $FAi$ is the intraindividual factor of asymmetry. Virtual *flies* with two eyes have been created by making all possible pairs of the 100 eyes.

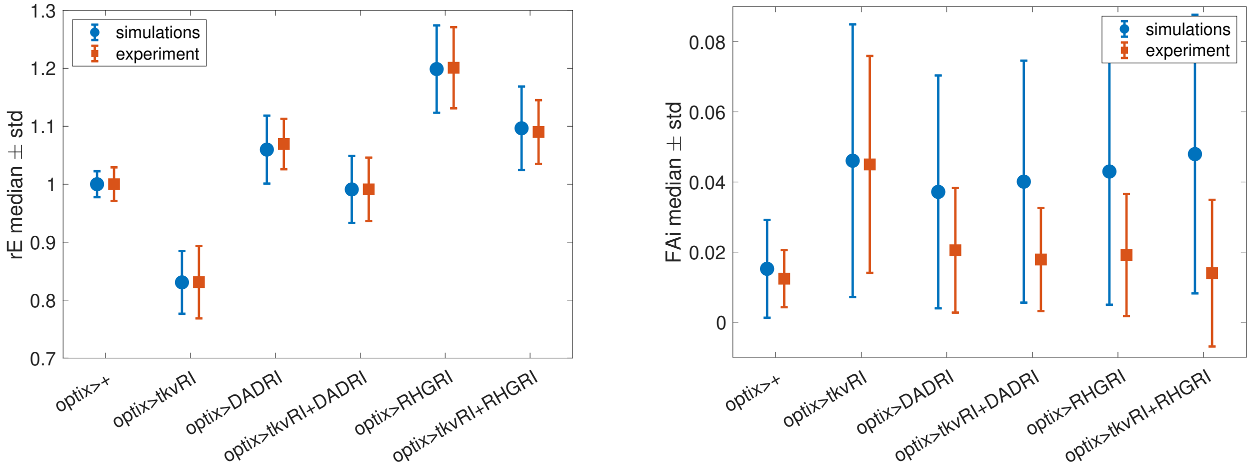


Figure 5: Weak apoptotic noise and strong proliferation noise: comparison for different conditions, as indicated, of simulation and experimental results. Simulations of 100 eyes with $V_a = 100$, $V_p = 20$, other parameters as in Tables 1–3. Left: eyes size, normalized to allow comparison of simulation and experiment. $rE$ is the average intraindividual eye size. Right: eyes asymmetry. $FAi$ is the intraindividual factor of asymmetry. Virtual *flies* with two eyes have been created by making all possible pairs of the 100 eyes.

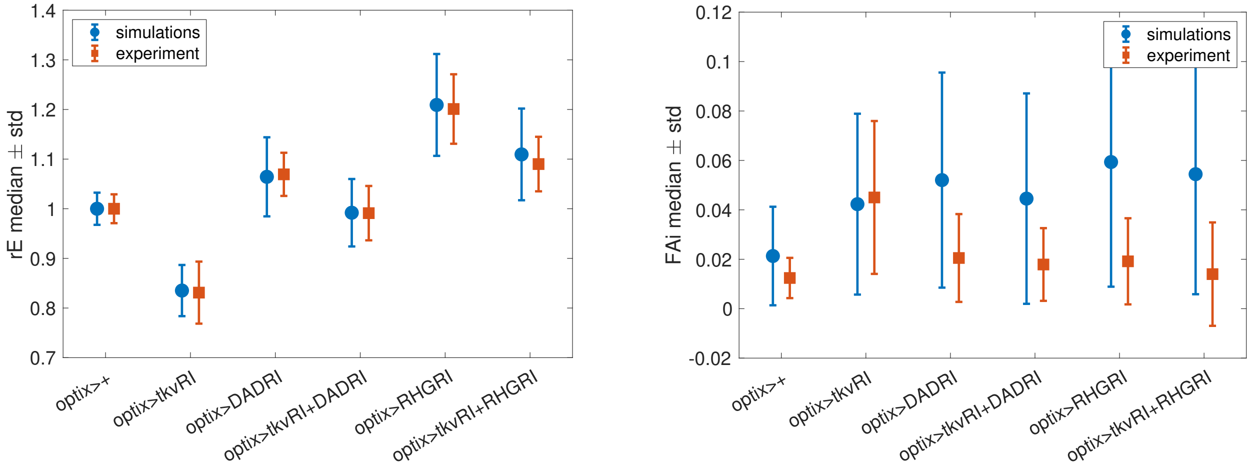


Figure 6: Weak apoptotic noise and strong initial differentiation noise: comparison for different conditions, as indicated, of simulation and experimental results. Simulations of 100 eyes with $V_a = 100$, $V_i = 15$, other parameters as in Tables 1–3. Left: eyes size, normalized to allow comparison of simulation and experiment. $rE$ is the average intraindividual eye size. Right: eyes asymmetry. $FAi$ is the intraindividual factor of asymmetry. Virtual *flies* with two eyes have been created by making all possible pairs of the 100 eyes.

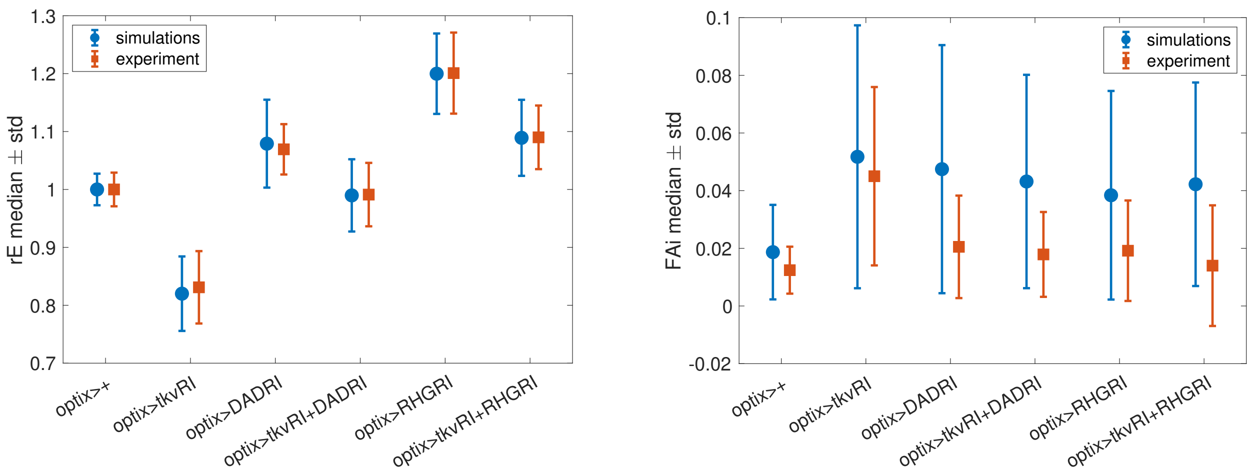


Figure 7: Weak apoptotic noise and strong final differentiation noise: comparison for different conditions, as indicated, of simulation and experimental results. Simulations of 100 eyes with $V_a = 100$, $V_f = 4$, other parameters as in Tables 1–3. Left: eyes size, normalized to allow comparison of simulation and experiment. $rE$ is the average intraindividual eye size. Right: eyes asymmetry. $FAi$ is the intraindividual factor of asymmetry. Virtual *flies* with two eyes have been created by making all possible pairs of the 100 eyes.

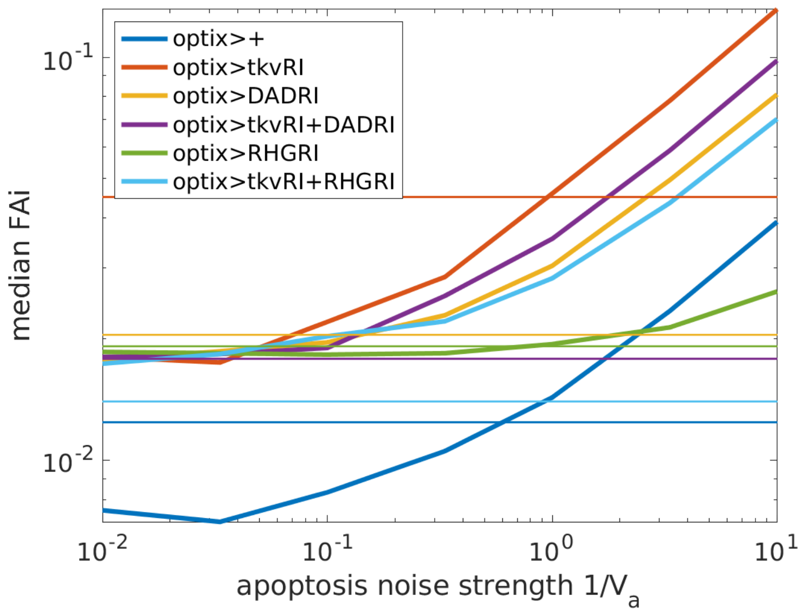


Figure 8: Fluctuating asymmetry index, $FAi$, of different conditions, as indicated, for a range of values of the apoptotic noise strength. Simulations of 1000 eyes, parameters as in Tables 1–3. Thin horizontal lines are $FAi$ experimental values for the corresponding condition. Virtual *flies* with two eyes have been created by making all possible pairs of the 200 eyes.

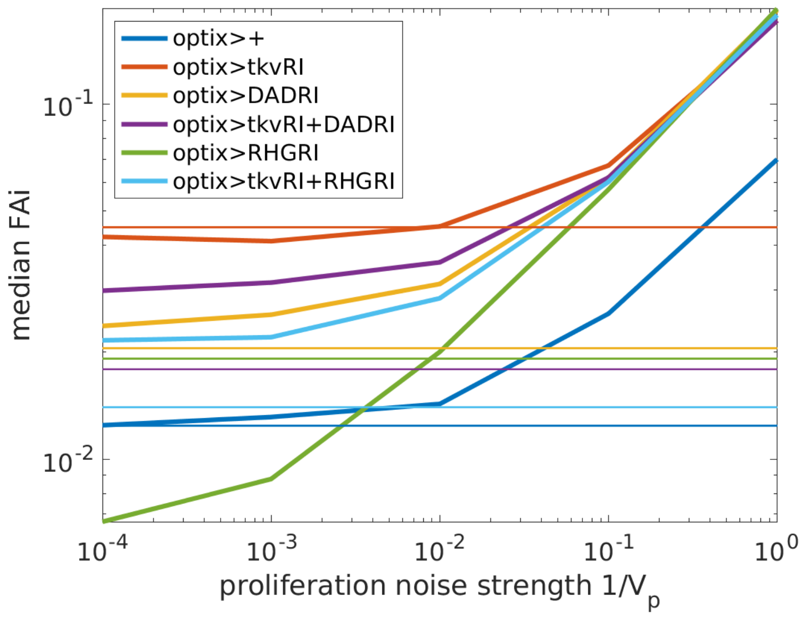


Figure 9: Fluctuating asymmetry index, $FAi$, of different conditions, as indicated, for a range of values of the proliferation noise strength. Simulations of 1000 eyes, parameters as in Tables 1–3. Thin horizontal lines are $FAi$ experimental values for the corresponding condition. Virtual *flies* with two eyes have been created by making all possible pairs of the 200 eyes.

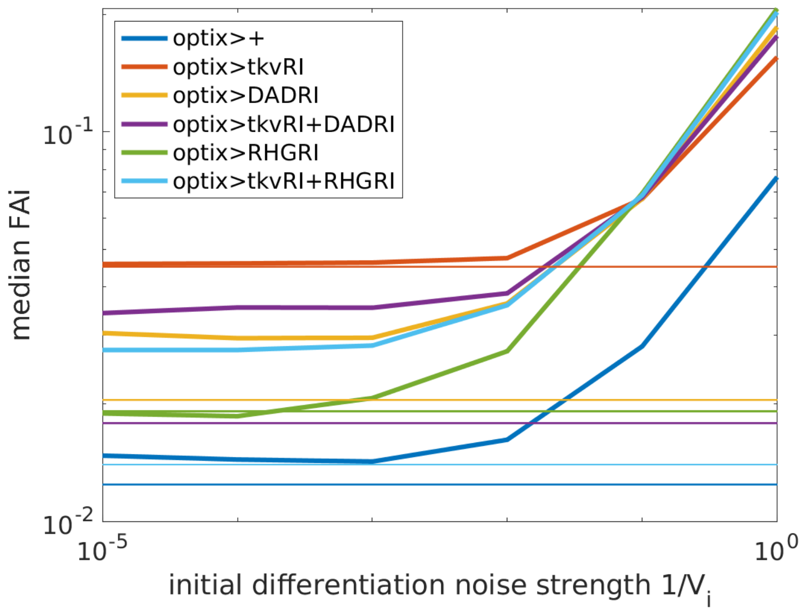


Figure 10: Fluctuating asymmetry index, $FAi$, of different conditions, as indicated, for a range of values of the initial differentiation noise strength. Simulations of 1000 eyes, parameters as in Tables 1–3. Thin horizontal lines are $FAi$ experimental values for the corresponding condition. Virtual *flies* with two eyes have been created by making all possible pairs of the 200 eyes.

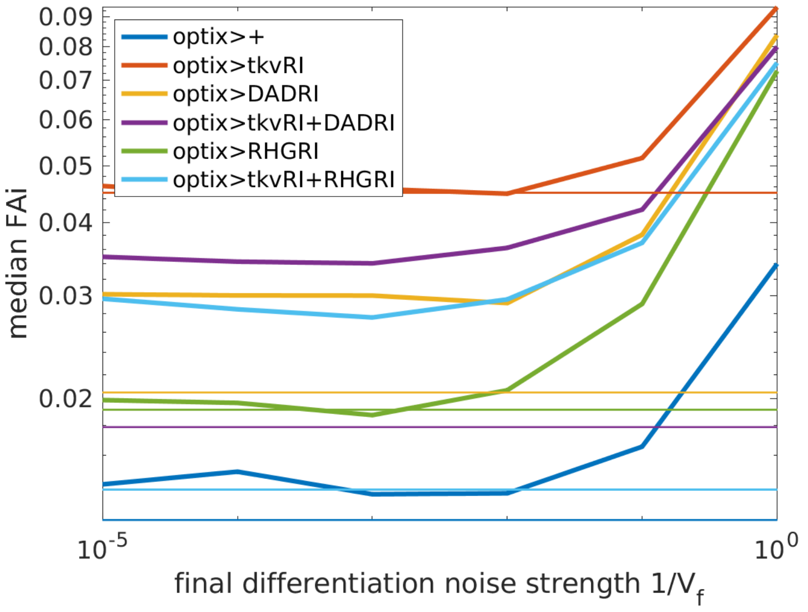


Figure 11: Fluctuating asymmetry index, $FAi$, of different conditions, as indicated, for a range of values of the final differentiation noise strength. Simulations of 1000 eyes, parameters as in Tables 1–3. Thin horizontal lines are $FAi$ experimental values for the corresponding condition. Virtual *flies* with two eyes have been created by making all possible pairs of the 200 eyes.

Asymmetry index, $FAi$, for a range of values of each of the noise strength parameters while keeping the others constant, Figures 8–11. In Figure 8 we see that for apoptotic noise strength the asymmetry is roughly the same in all conditions (please recall that for *optix>+* we have introduced a noise reducing factor). For high enough apoptotic noise the condition with enhanced apoptosis, *optix>tkvRI*, systematically presents higher asymmetry than any other condition (and please observe that the vertical axis is logarithmic). Figures 9–11 show that, once the apoptotic noise rate $V_a$ is fixed at a value producing strong noise, low noise strengths of the other mechanisms are highly irrelevant for setting the values of $FAi$. Once those noises become strong enough, they tend to make all conditions equally asymmetric (recall again the noise reducing factor for *optix>+*), therefore showcasing that strong apoptotic noise is necessary for the qualitative observation of high asymmetry in the high apoptosis *optix>tkvRI* condition.

#### SENSITIVITY ANALYSIS

To asses the robustness of our results to parameter values, we have performed a sensitivity analysis [13, 14]. In our study, we characterize the sensitivity $S_{YX}$ of the measurable variable $Y$ in relation to variations in the parameter $X$ as:

$$S_{YX} = \frac{\partial \log Y}{\partial \log X} \tag{35}$$

This metric is assessed at a specific location within the parameter domain. The logarithmic differ-

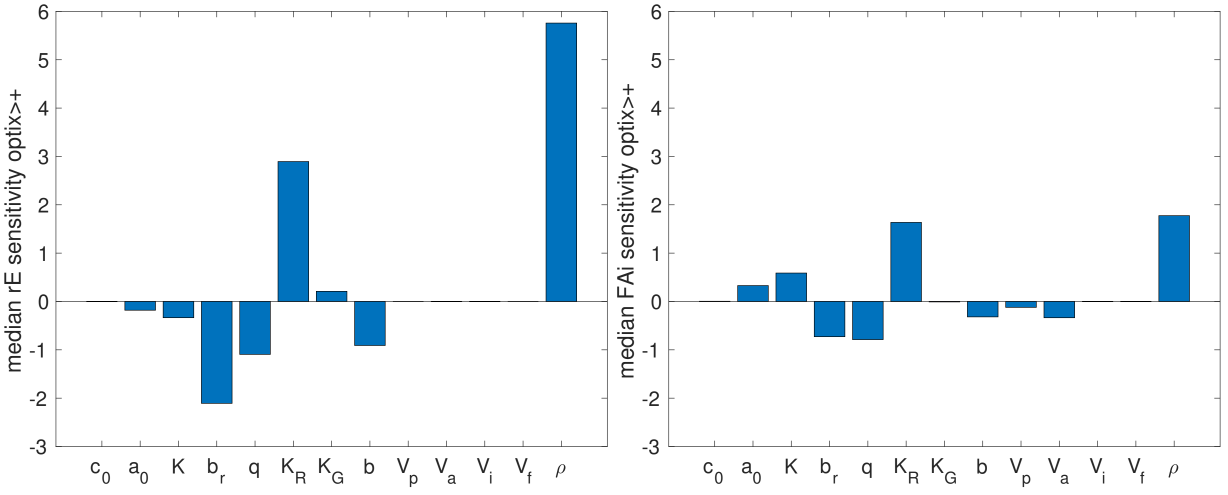


Figure 12: Sensitivity of the medians of $rE$ (left panel) and $FAi$ (right panel), calculated from 1000 realizations, with respect to 10% changes in the indicated parameter values. This analysis is computed at the point in parameter space used to describe the *optix>+* condition, Tables 1 and 3. Analysis of other conditions show similar results.

ential outlined above can be reformulated as:

$$S_{YX} = \frac{dY/Y}{dX/X} \tag{36}$$

For minor fluctuations in the parameter $X$, Eq. (36) can be approximated by the proportion of relative changes, expressed as:

$$S_{YX} = \frac{\Delta Y/Y}{\Delta X/X} \tag{37}$$

The model's sensitivity to shifts in the parameter $X$ is ascertained by examining the chosen variable $Y$ at two distinct points: the reference point $X_0$ and $X = X_0 + \Delta X$. Therefore, $S_{YX}$ is defined as the quotient of the percentual variation in $Y$ to the percentual variation in $X$. To determine senistivity values, we have used values of $\Delta X$ corresponding to 10% of the value of $X_0$. Since our model is stochastic, we have run 1000 simulations and determined the sensitivities of the medians of $rE$, the average intraindividual eye size, and $FAi$, the intraindividual factor of asymmetry. The results for medians of 100 simulations are virtually indistinguishable, showing that the caluclation has converged.

Figure 12 shows the results of our sensitivity analysis. Together to the parameters in Tables 1 and 3, we were interested in calculating the sensitivity to the proliferation rate. Since we had defined the timescales of all paramaters with respect to it, we have to introduce an explicit parameter that only affects proliferation rate. This is the parameter $\rho$ in the following version of the drift equation of our model:

$$F(X) = \frac{\log(2)}{c_0 R} \begin{pmatrix} \rho(1 - a(R_n))G - a(R_n)G - f(G)h(R) \\ f(G)h(R) - b\,h(R)R_n \\ b\,h(R)R_n \end{pmatrix}. \tag{38}$$

This non-dimensional parameter has a value of $\rho = 1$, therefore the sensitivity analysis is donde comparing it to the results with a 10% change, corresponding to a value of $\rho' = 1.1$. Unsurprisingly, we find that this is the parameter to which the eye size is most sensitive. However, the intraindividual factor of asymmetry is much less sensitive to changes in growth rate: a trend for bigger eyes naturally allows for larger variations, but the resulting fluctuations in eye size are much constrained. This is a general property: Figure 12 shows that our model is robust to parameter values, and for those parameters with a higher sensitivity of the eye size, namely the proliferation rate $\rho$ and the parameters $K_R$ and $b_r$ defining the initial differentiation rate, the corresponding sensitivity of the assymetry factor is sensibly lower. In particular, the model is very insensitive to small variations in the parameters determining noise strength, showing that our conclusions are in not dependent on a fine tuning of these parameters. The effects of large variations in the noise parameters was discussed in the previous section.
